# Bayesian semi-nonnegative matrix tri-factorization to identify pathways associated with cancer phenotypes

**DOI:** 10.1101/739110

**Authors:** Sunho Park, Nabhonil Kar, Jae-Ho Cheong, Tae Hyun Hwang

## Abstract

Accurate identification of pathways associated with cancer phenotypes (e.g., cancer sub-types and treatment outcome) could lead to discovering reliable prognostic and/or predictive biomarkers for better patients stratification and treatment guidance. In our previous work, we have shown that non-negative matrix tri-factorization (NMTF) can be successfully applied to identify pathways associated with specific cancer types or disease classes as a prognostic and predictive biomarker. However, one key limitation of non-negative factorization methods, including various non-negative bi-factorization methods, is their lack of ability to handle non-negative input data. For example, many molecular data that consist of real-values containing both positive and negative values (e.g., normalized/log transformed gene expression data where negative value represents down-regulated expression of genes) are not suitable input for these algorithms. In addition, most previous methods provide just a single point estimate and hence cannot deal with uncertainty effectively.

To address these limitations, we propose a Bayesian semi-nonnegative matrix tri-factorization method to identify pathways associated with cancer phenotypes from a real-valued input matrix, e.g., gene expression values. Motivated by semi-nonnegative factorization, we allow one of the factor matrices, the centroid matrix, to be real-valued so that each centroid can express either the up- or down-regulation of the member genes in a pathway. In addition, we place structured spike-and-slab priors (which are encoded with the pathways and a gene-gene interaction (GGI) network) on the centroid matrix so that even a set of genes that is not initially contained in the pathways (due to the incompleteness of the current pathway database) can be involved in the factorization in a stochastic way specifically, if those genes are connected to the member genes of the pathways on the GGI network. We also present update rules for the posterior distributions in the framework of variational inference. As a full Bayesian method, our proposed method has several advantages over the current NMTF methods which are demonstrated using synthetic datasets in experiments. Using the The Cancer Genome Atlas (TCGA) gastric cancer and metastatic gastric cancer immunotherapy clinical-trial datasets, we show that our method could identify biologically and clinically relevant pathways associated with the molecular sub-types and immunotherapy response, respectively. Finally, we show that those pathways identified by the proposed method could be used as prognostic biomarkers to stratify patients with distinct survival outcome in two independent validation datasets. Additional information and codes can be found at https://github.com/parks-cs-ccf/BayesianSNMTF.

## 1. Introduction

Accurate identification of pathways associated with cancer phenotypes (e.g., cancer sub-types and treatment outcome) enables us to understand better molecular biology processes in cancer and could lead to discovering reliable prognostic and/or predictive biomarkers for better patients stratification and treatment guidance. Non-negative matrix tri-factorization (NMTF) models can provide an intuitive and efficient way to identify associations between two different entities by simultaneously clustering rows and columns of the data matrix.^1^ In our previous work^2^ (referred to as NTriPath), we use NMTF to identify pathways associated with cancer types from mutation data: the mutation data matrix is decomposed into the cancertype indicator matrix, the association matrix between cancer types and pathways, and the centroid matrix (each centroid corresponds to the pattern of gene mutations within each pathway). Pathway membership information, e.g., gene-pathway annotations from Kegg pathway database, and a gene-gene interaction (GGI) network are incorporated into the factorization model through the framework of regularized optimization. It is shown from the The Cancer Genome Atlas (TCGA) data that the top pathways ranked by the method are closely related to clinical outcomes.^2^ However, this approach has several limitations. First, the input matrix is restricted to be non-negative and hence cannot readily model many types of genomic data, including copy number alteration and normalized/log transformed gene expressions, which are real-valued. Second, the method provides just a single point estimate of the model’s parameters and thus cannot deal with uncertainty well. Moreover, it involves many hyper-parameters, e.g., regularization constants, which should be tuned carefully. However, since the association identification from the input (mutation) matrix is clearly an unsupervised problem, i.e., there is no corresponding output for the input matrix, it is not clear how to find the optimal hyperparameter values for the given input data.

To address the aforementioned limitations of NTriPath, we propose a novel Bayesian semi-nonnegative matrix factorization model, where the biological prior knowledge represented by a pathway database and a GGI network is incorporated into the factorization through structured spike-and-slab sparse priors.^3^ First, in order to handle real-valued input data, e.g., gene expression values, we allow one of the latent (factor) matrices, the centroid matrix, to have positive and negative values so that each centroid (corresponding to a pathway) can express the up-regulation or the down-regulations of the member genes in the pathway. Second, we encode pathway membership information and a GGI network into the factorization model through the framework of Bayesian learning. Specifically, we model the priors over the centroid matrix matrix using the structured spike-and-slab distributions, where our prior knowledge of the sparsity pattern is encoded into the prior distributions thorough underlying Gaussian processes (GPs).^3^ To conclude the prior modeling for the centroid matrix, we define the mean vectors and covaraince matrices of the GPs using the pathway membership information and the GGI network. As a result, even non-member genes of the pathways can be involved in the factorization in a stochastic manner. Note that our method is a full Bayesian approach: priors are placed on the model’s parameters (the latent matrices) and hyper-parameters (e.g., the noise precision) and updated by observations (resulting in the posteriors). Thus, in contrast to NTriPath, which relies on only the single most probable setting of the model’s parameters and hyper-parameters (regularization constants), our method produces more robust factorization results by averaging over all possible settings. Finally, we propose the update rules for the posterior distributions by utilizing the framework of variational inference. Using experiments on synthetic datasets, we show the superiority of our proposed method over NTriPath (where a folding approach^4^ is used to deal with negative values in the input matrix). Using TCGA gastric cancer and metastatic gastric cancer immunotherapy clinical-trial datasets,^5^ we show that the proposed method could identify biologically and clinically-relevant pathways associated with TCGA gastric cancer molecular sub-types and immunotherapy response. Finally we show that those pathways identified by our method could be used as prognostic biomarkers to stratify patients with distinct survival outcome in two independent validation datasets.

### Notations

For a matrix ***A***, ***a_i_*** represents its ith row vector, i.e., (***A_i_***,:)^T^. Similarly, 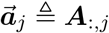 refers to its *j*th column vector. The (*i, j*)th element of the matrix ***A*** is expressed by *A_ij_*.

## 2. Background

Non-negative matrix factorization (NMF), which here refers to the matrix bi-factorization (decomposing a matrix into two smaller matrices), has been applied to many different biological problems as a tool for clustering, dimensionality reduction and visualization (please see references herein^6^). It provides a parts-based local representation, making NMF unique compare to other linear dimensionality reduction methods such as principal component analysis (PCA). However, NMF is limited to non-negative input data. When the input matrix contains positive and negative values, a natural way is to decompose the input matrix into a centroid matrix (assumed to be real-valued) and a cluster membership indicator matrix (assumed to be non-negative). This approach is the main motivation of semi-nonnegative factorization,^7^ and we use this same idea to allow our method to find patterns from real-valued input data.

The spike-and-slab prior is the standard approach for sparse learning, which is the selection of a subset of features from high-dimensional input data. It can be expressed as a mixture of a point mass at 0 (spike) and a continuous distribution (slab):

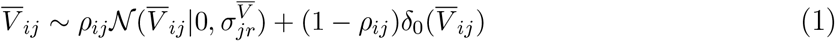

where 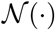 is a Gaussian distribution, *ρ_ij_* ∈ [0,1] is a mixing coefficient, and *δ*_0_(·) is Dirac delta funciton, i.e., 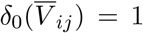 at 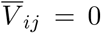, and 0 elsewhere. The mixture structure of the spike-and-slab prior can produce a bi-separation effect where the posterior distributions over the coefficients for *irrelevant* features are peaked at zero while those over the coefficients of *relevant* features have a large probability of being non-zero. The spike-and-slab prior (1) can be equivalently rewritten with a binary variable, and the posterior mean of this binary variable indicates how the corresponding coefficient is actually different from zero.

## 3. Bayesian Semi-Nonnegative Tri-Matrix Factorization (Bayesian SNTMF)

We propose a Bayesian method to identify associations between cancer phenotypes (e.g., molecular subtypes) and pathways from human cancer genomic data. In this work, we consider only gene expression data, but our method can be applied to other data types that can be formed into real-valued matrices, e.g., copy number and miRNA expression. We develop a semi-nonnegative matrix tri-factorization method in the framework of Bayesian learning, where the prior knowledge represented by a pathway membership information and a GGI network is taken into account in the factorization through structured spike and slab prior distributions.^3^

### 3.1. Model formulation

We assume that observations are given in the form of a matrix ***X*** ∈ ℝ^*N×D*^ where *X_ij_* represents the ith patient’s expression value for the *j*th gene, and *N* and *D* are the number of samples and genes, respectively. We assume that pathway information is also given in a form of a matrix ***Z***^0^ ∈ ℝ^*D×R*^ where each element represents the membership of a gene to a pathway, i.e., 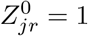 if the *j*th gene is a member of the *r*th pathway, and 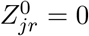 otherwise. Our main objective is to approximate ***X*** as a product of three latent matrices added with residuals ***E*** ∈ ℝ^*N×D*^:

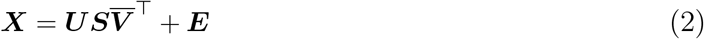

where 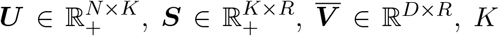 is the number of the sub-types, and *R* is the number of the pathways. We assume that the matrix ***U*** is constructed from patient clinical data: *K* is the number of sub-types we are interested in, and *U_ij_* = 1 indicates that the *i*th patient is of the *j*th sub-type (1-of-K encoding, i.e., *U_ik_* ∈ {0,1} and 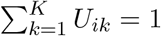). The realvalued matrix 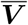 consists of *R* basis vectors, the rth column of which is a pattern associated with a corresponding pathway: only few elements (corresponding to the member genes of a pathway, i.e., 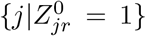) would have non-zero values, representing either over-expression 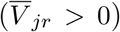 or under-expression 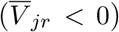, and all other elements are set to zero. Then, the non-negative matrix ***S*** encodes associations between the sub-types and the pathways, where each element *S_ij_* represents the association between the *i*th sub-type and the *j*th pathway. Once ***S*** is learned, we can easily identify pathways related to a certain sub-type by selecting the top pathways that have the largest values in the corresponding row in S. As all the latent variables are learned in the Bayesian learning framework, the likelihood of the model and the prior distribution over the latent variables are defined according to our model assumptions.

Assuming the residuals *E_ij_* in eq. (2) to be sampled from i.i.d. Gaussian distributions with mean zero and precision *γ*, we can specify the likelihood of the factorization model:

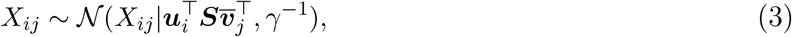

where the precision *γ* (the inverse of the variance) is sampled from a Gamma distribution.

The following discusses how we define the priors over the latent variables. For ***S***, each element is assumed to be sampled from an Exponential distribution to ensure its non-negativity:

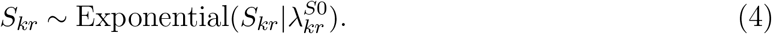

For 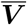, the simplest inference approach would be to calculate the posterior distributions (with Gaussian distribution priors) over only the elements in the matrix that are corresponding to the member genes in the pathways, i.e., 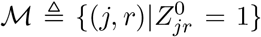, and leave the other elements as zero. However, it is widely accepted that pathway databases are not complete, that there are unknown missing genes in a pathway. To include unknown missing member genes in the pathways into the factorization, we use the concept of sparse learning, where sparse prior distributions (e.g., spike-and-slab or Laplace distributions) are placed over all the elements of 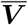 and only few elements (including those in the set 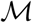) are encouraged to have non-zero values. We make use of a gene-gene interaction network as well as of the pathway information ***Z***^0^ to determine the the positions of non-zero elements in 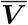 based on the assumption that two connected genes in the graph would more likely to be active together in a pathway. Denote a gene-gene interaction network by ***A*** ∈ ℝ^*D×D*^, where *A_jj′_* = 1 if genes *j* and *j*′ are connected on the network, and *A_jj′_* = 0 otherwise, and assume that there is no self connection, i.e., *A_jj_* = 0. We then will show that the priors incorporating ***Z***^0^ and ***A*** can be defined using the structured spike and slab prior model^3^ which imposes spatial constraints on spike-and-slab probabilities through a Gaussian process (GP). We define a GP for each pathway and encode the mean vector and covariance matrix of the GP using our prior knowledge given by ***Z***^0^ and ***A***.

With reparametrization of the variable 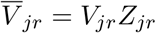 (*Z_jr_* is assumed to be a binary variable, i. e., *Z_jr_* ∈ {0,1}), where 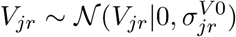) and *Z_jr_* ~ Bernoulli(*ρ_jr_*), the spike-and-slab prior over 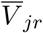 in (1) can be equivalently written for the new variables *V_jr_* and *Z_jr_*:

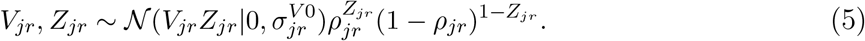

We can consider the binary variable *Z_jr_* as a on-off switch which determines whether *V_jr_* is included into the factorization model. To connect ***Z***^0^ and ***A*** to *Z_jr_*, we define the parameter of the Bernoulli distribution *ρ_jr_* in the following hierarchical way based on the frame of GP:

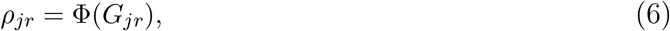

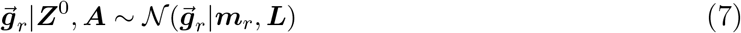

where 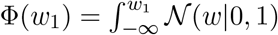 is a cumulative standard Gaussian distribution function and 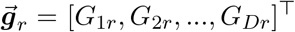. Each element of the mean vector ***m**_r_* is set according to the membership information encoded in ***Z***^0^: *m_jr_* = *ξ*_+_ where *ξ*_+_ > 0 if 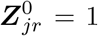, and *m_jr_* = *ξ*_−_ where *ξ*_−_ < 0 otherwise (the more negative value *ξ*_−_ is, the more sparse prior we get). The covariance matrix ***L*** is set to a normalized Laplacian matrix ***L*** = ***I*** — ***D***^-1/2^ ***AD***^-1/2^, where ***D*** is a diagonal matrix whose ith diagonal element is a summation of the *i*th row of the matrix ***A***. Combining all these assumptions, we can see that if gene *i* (a nonmember of the rth pathway) has connections to the member genes on the network, then *G_ir_* would become high and its on-off binary variable *Z_ir_* is more likely to be one. Note that 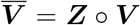. The binary matrix ***Z*** is determined by a stochastic process, and thus the elements in ***V*** that even are not in the set 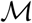 (originally not in the pathways) can contribute to the factorization model.

As a result, our factorization model can be summarized as follows:

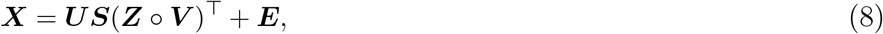

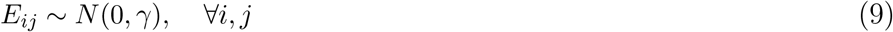

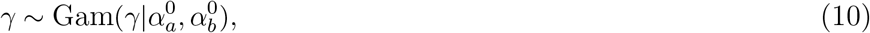

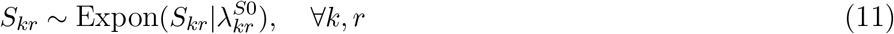

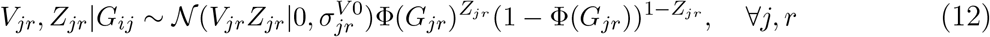

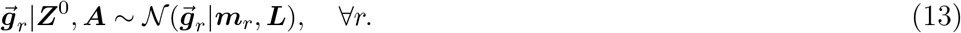

The conceptual view of our method is depicted in Figure 1.

**Fig. 1.**
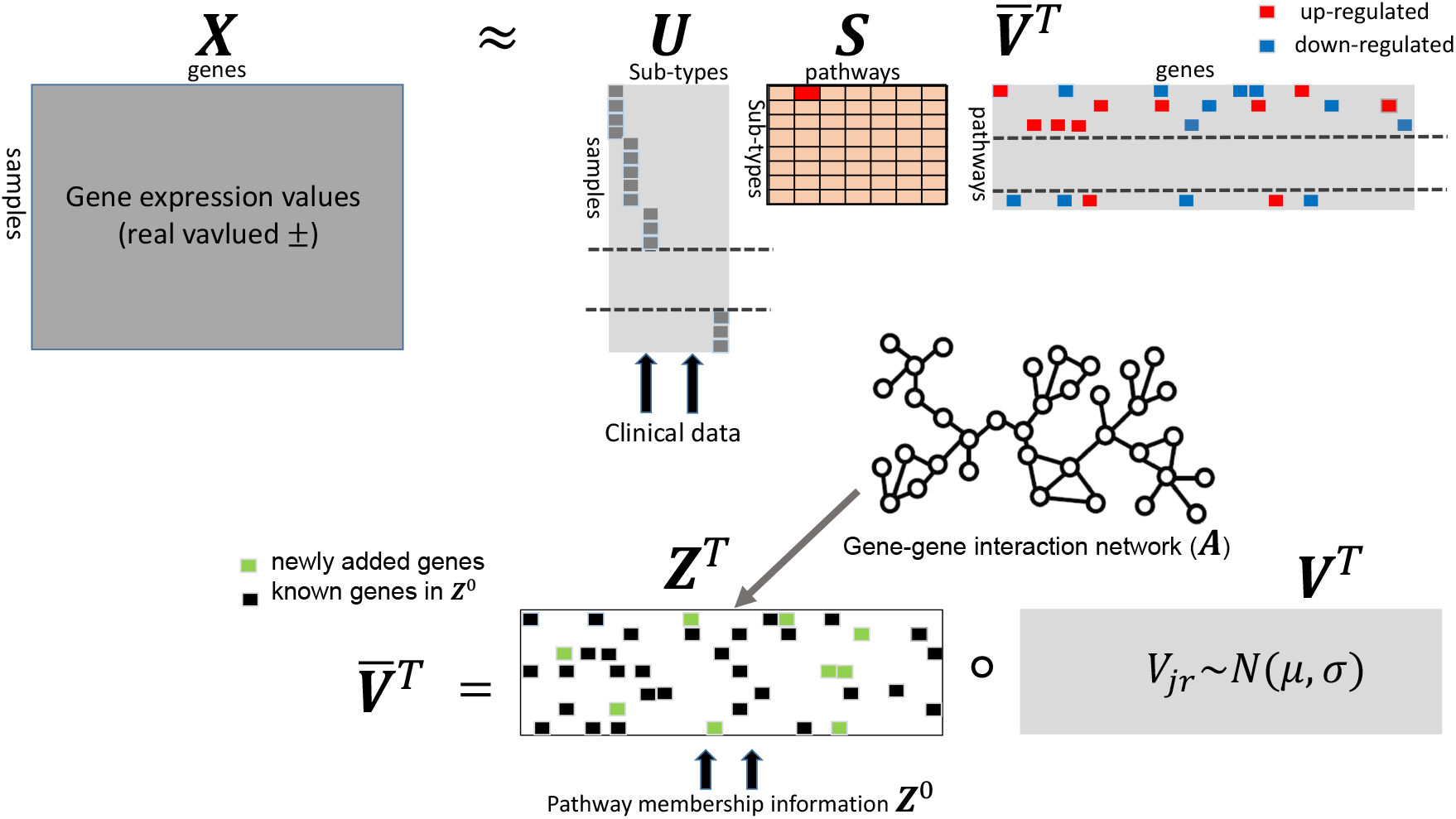
The input matrix is decomposed into ***U*** (samples × sub-types), ***S*** (sub-types × pathways), and 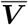 (genes × pathways). The centroid matrix 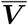 is further decomposed into the binary indicator matrix ***Z*** and the genome-wide pattern matrix ***V***. We encode the pathway membership information ***Z***^0^ and the GGI network ***A*** into the binary matrix ***Z*** through the structure spike-and-slab priors.

### 3.2. Variational inference

We approximate the posterior distributions over all the latent variables in the variational inference framework as their close form expressions are not available. We assume that the variational distributions are factorized as follows:

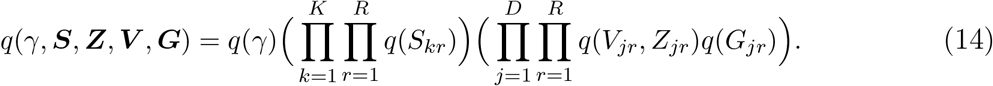

Note that the elements in the latent matrices (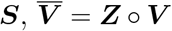, and ***G***) are assumed to be fully factorized. The form of each variational distribution is assumed to be as follows

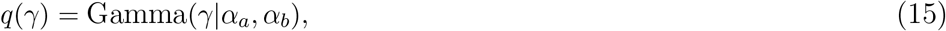

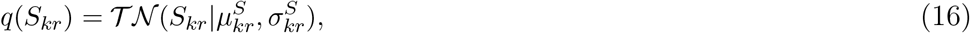

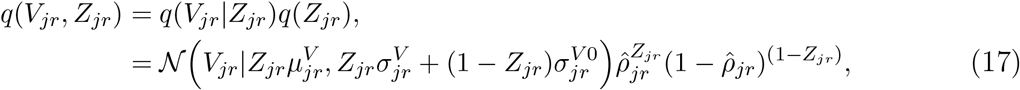

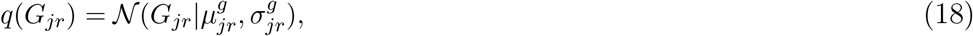

where 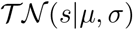 represents a truncated Normal distribution defined on the nonnegative region *s* ≥ 0, i.e., 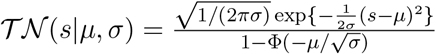 if *s* ≥ 0, and 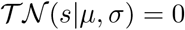 otherwise. Denoting a set of all the latent variables by Θ = {*γ*, ***S, Z, V, G***}, the variational distribution, *q*(Θ), can be obtained by maximizing the variational lower bound with respect to *q*(Θ):^8^

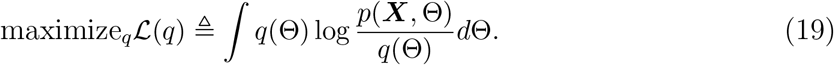

Note that the variational bound 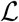 is a lower bound on the log-likelihood, i.e., 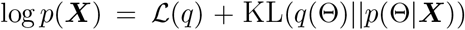, where the second term in RHS is the Kullback-Leibler (KL) divergence between the variational distribution and the true posterior distribution and always nonnegative. Thus, finding the optimal variational distributions by solving the optimization problem (19) can be easily justified. For each step, we update one variational distribution, fixing the others, and we then proceed to cyclically update all variational distributions in this manner. Based on combining the inference methods for Bayesian non-negative matrix tri-factorziation in^9^ and for spike-and-slab prior distributions, the variational distributions *q*(*γ*), {*q*(*S_kr_*)} and {*q*(*V_jr_, Z_jr_*)}, can be updated in closed form. For {*q*(*G_jr_*)}, their means and variances can be updated by any iterative gradient-based optimization methods, e.g., limited-memory BFGS used in our experiments. More detailed derivations are found in our supplementary material available at https://github.com/parks-cs-ccf/BayesianSNMTF.

## 4. Experimental results

We conduct experiments on both simulation and real-world datasets: 1) using the simulation datasets, we show how our method works and display the superiority of our method over NtriPath (which is a point estimate method); 2) using the two gastric cancer datasets, we demonstrate that the our method can identify biologically and clinically-relevant pathways associated with the molecular sub-types in gastric cancer as well as immunotherapy response and validate these results on independent validation datasets.

We here discuss how to find pathways closely associated with each sub-type based on the factorization results from our method, as the final outputs of our method are the variational distributions (the approximate posteriors) over the latent variables, including the association matrix ***S***. Specifically, we simply use the posterior mean of each variable as its estimate. We denote the estimate of each latent matrix *M* by 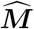, where each element represents the posterior mean of the corresponding element in the matrix *M* (please refer to our supplementary material to see how to calculate the mean value of each posterior distribution). For the estimate association matrix 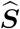, which is always non-negative, we can easily see that the larger 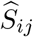 is, the stronger association between the *i*th sub-type and the *j*th pathway. Lastly, we explain how to initialize some variables in our model. For the mean vectors of the GPs (***G***), we set *ξ*_+_ = 5 and *ξ*_−_ = –5 for all the experiments, which means that we assume a strong prior belief on the initial pathway information ***Z***^0^. However, as we will see from the experiment with simulation datasets, our method is able to recover missing pathway membership. The detailed information on the initialization for our method is included in the supplementary material.

### 4.1. Simulation datasets

With this simple example, we first show how our method works in the case of incomplete pathway membership information. We generate the observation matrix ***X*** ∈ ℝ^300×400^, where the matrix contains 3 sub-types and each sub-type shows a unique pattern, one or two blocks of up- or down-regulated genes in each sub-type (***X*** in Figure 2-(a)). Elements in the pattern blocks are drawn from either 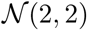 for the up-regulation case or 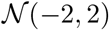 for the down-regulation case, but elements in the non-pattern blocks are assumed to be background noise and are sampled from 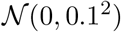. We construct the sub-type indicator matrix ***U*** based on our knowledge on the sub-type information. We generate a pathway membership matrix ***Z***_0_ according to the block structure of the input matrix ***X*** such that the true associations between the sub-types and the pathways can be easily identifiable (***Z***^0*T*^ in Figure 2 (b)). Note that we assume the pathway membership matrix ***Z***^0^ incomplete: we randomly remove 80% of member genes from one of the blocks in the 3rd pathway. For the gene-gene interaction network, we randomly connect two genes on the network with probability 0.1.

**Fig. 2.**
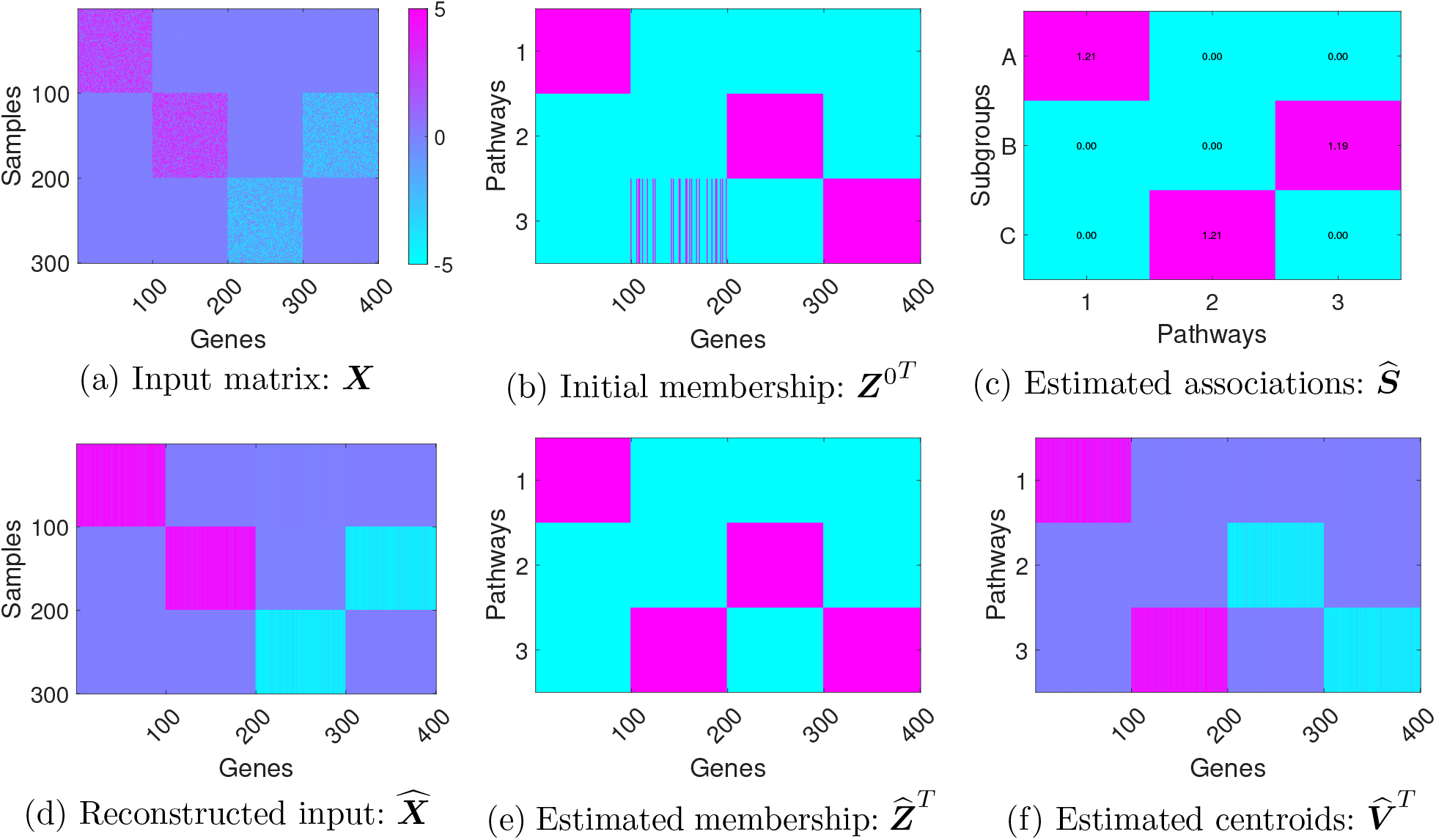
Factorization results of the simulation data under the assumption that the pathway membership information might be incomplete: multiple member genes in one of the pathways are missed (b). The results indicate that our method can successfully recover the membership information (e).

Figure 2 (c)-(f) shows that our factorization method works well even with the incomplete pathway information. Figure 2 (c) indicates that our method can accurately estimate true associations between sub-types and pathways. For example, the pathway associated with the 2nd sub-type (which includes the samples 101 to 200 in the input data) is the 3rd pathway as we designed, and we can easily confirm this association from the estimate association matrix 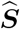 because only 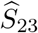 has a significantly high value and the others, 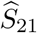 and 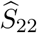, are zero. This result is the same for the other sub-types. we also see that our method can successfully recover the pathway membership information from the data (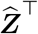 in Figure 2 (e)). This is a promising result considering current pathway databases might be incomplete as our knowledge on molecular biology processes is incomplete. Finally, we can see that our method can correctly find the up/down regulation patterns from the real-valued input data (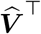 in Figure 2 (f)).

We also test our method on an additional simulation dataset to show the superiority of our method over NTriPath. For non-negative factorization methods, one of standard ways to deal with negative values in the input matrix is to fold the matrix by columns:^4^ every column will be represented in two new columns in a new matrix, one of which contains only positive values and the other only the magnitudes of negative values. This approach doubles the number of columns in the original matrix and thus causes additional computational burdens, e.g., the GGI network becomes 2^2^ times larger. Moreover, it breaks the original patterns in the input matrix because non-negative and negative values are separately processed. In addition, we can see that our method is more robust against noise in general than NTriPath, as Bayesian methods deal with uncertainty more effectively than point estimate methods which rely on a single most probable setting of the model’s parameters. Detailed information about this experiment is included in our supplementary material.

### 4.2. TCGA gastric cancer and metastatic gastric cancer immunotherapy clinical-trial datasets

We first identify the top pathways associated with: 1) molecular sub-types in the TCGA gastric cancer (GC) data; and 2) response/non-response in the metastatic gastric cancer (mGC) immunotherapy clinical-trial data.^5^ We then validate the pathways identified by our method in both datasets by investigating if these pathways could be used as prognostic biomarkers to stratify patients from two validation datasets, ACRG^10^ and MDACC,^11^ into groups with distinct survival outcome.

We provide brief descriptions of the datasets with the notations used in Section 3. For TCGA GC data (*N* = 277), we download the normalized gene expression (mRNA) data^*^. The samples are divided into *K* = 4 groups according to their molecular sub-types: Epstein-Barr virus (EBV), microsatellite instability (MSI), genomically stable (GS), and chromosomal instability (CIN). For the immunotherapy response for mGC data (*N* = 45), we download the gene expression data from,^5^ which is normalized by FPKM, and additionally apply log-transformation and standardization. The data includes the patients’ treatment outcomes, which are categorized into 4 sub-types: complete response (CR), partial response (PR), progressive disease (PD), and stable disease (SD). In order to find more distinguishable patterns between groups, we here divide the samples into just *K* = 2 groups: responders (CR+PR) and non-responders (PD+SD). Next, we download a GGI network (***A***) from ^†^ and use *R* = 4,620 sub-networks from^12^ to define the pathway membership matrix ***Z***^0^. After combining all these different data sources, the numbers of the input genes are *D*_1_ = 14, 787 and *D*_2_ = 15,347 for TCGA gastric cancer data and the immunotherapy response data, respectively. The information of both datasets is summarized in Table 1.

**Table 1.** Summary of the two datasets, TCGA GC and mGC datasts.

| data | $N$ | $D$ | $K$ | phenotypes |
| --- | --- | --- | --- | --- |
| TCGA gastric cancer | 277 | 14,787 | 4 | {CIN vs EBV vs GS vs MSI} |
| Immunotherapy response | 45 | 15,347 | 2 | {responder vs non-responder} |

After training our factorization model on each dataset, we select the top 3 ranked path ways for each subtype based on the estimated association matrix 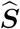 (12 = 3 × 4 pathways consisted of 83 genes are selected for TCGA GC data, and 6 = 3 × 2 pathways consisted of 36 genes for the immunotherapy response for mGC data). To assess biological relevance of identified top pathways from TCGA GC and immunotherapy for mGC datasets, we performed gene set enrichment analysis using PANTHER (http://www.pantherdb.org). We found that pathways identified by our method are enriched with biologically relevant pathways that are associated with cancer phenotypes. For example, 36 genes from mGC immunotherapy response data are enriched with positive regulation of TGFbeta pathway, T-cell migration, etc. Specifically, member genes of 36 gene signatures such as FN1 and FBLN1, involved with TGFbeta regulation are down-regulated and CCL5, CCL21, and CXCL13 which are involved with T-cell migration are up-regulated in response group compared to non-response group, respectively. Activation of TGFbeta pathway serves as central mechanisms to suppress immune system thus deactivation of TGFbeta may increase response to immunotherapy.^13^ Active T-cell migration into tumor microenvironment could increase response rates to immunotherapy and increase survival.^14^ These indicate that our proposed method utilizing real-valued input data could successfully identify down and/or up-regulated pathways that are biologically relevant to and associated with immunotherapy response. It is worth to note that these findings were not reported in the original work.^5^ Further details of pathway analysis are available at https://github.com/parks-cs-ccf/BayesianSNMTF.

To evaluate prognostic utility of 83 and 36 genes in the top 3 pathways from TCGA GC and mGC immunotherapy datasets, we perform a consensus clustering to stratify gastric cancer patients using two validation cohorts ACRG *(N* = 300) and MDACC *(N* = 267), respectively. Setting the number of clusters to 4, we run a consensus clustering method (500 NMF repetition with bootstrapping^15^) on gene expression values of the selected genes in each dataset and generate Kaplan-Meier (KM) plots using overall survival. We specifically choose the number of clusters as 4 in order to show whether our biomakers have comparative prognostic performance to TCGA GC molecular 4 sub-types, i.e., EBV, MSI, GS, and CIN. Figure 3 shows that subgroups identified by 83 and 36 genes from TCGA GC and mGC immunotherapy datasets have distinct survival outcomes which suggests that the pathways identified by our method can be served as prognostic biomarkers to stratify GC patients.

**Fig. 3.**
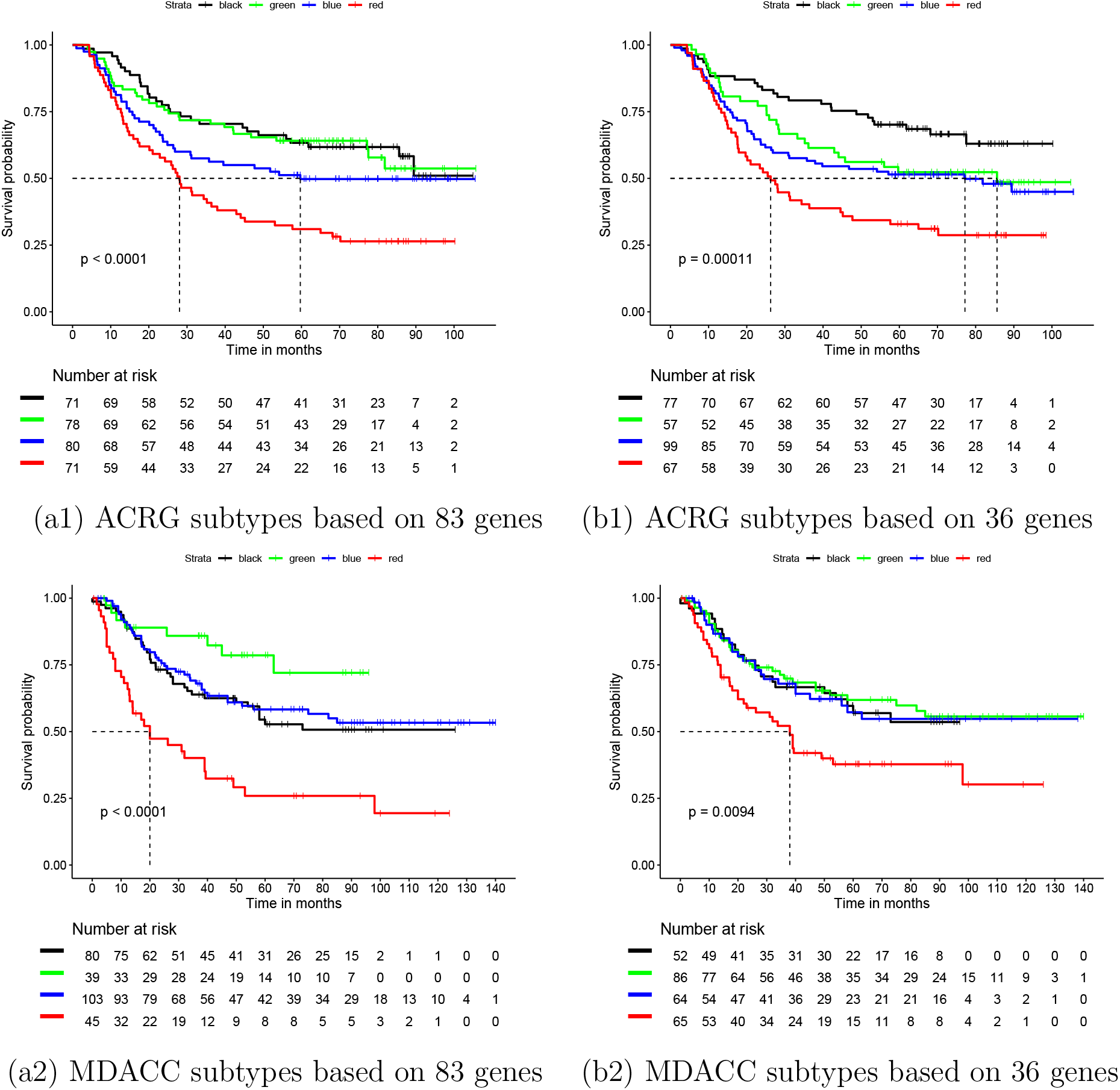
KM plots from ACRG and MDACC cohorts. In each of ACRG and MDACC validation cohorts, four subgroups clustered based on gene expression values of the 83 and 36 gene signatures from TCGA GC and the mGC immunotherapy response datasets, respectively. KM plots with log-rank test indicate that the subgroups identified by the 83 and 36 gene signatures have statistically significant different survival outcomes.

## 5. Discussion

We have proposed a Bayesian semi-nonnegative matrix tri-factorization method to identify associations between cancer phenotypes e.g., molecular sub-types or immunotherapy response, and pathways from the real-valued input matrix, e.g., gene expressions. Motivated by semi-nonnegative factorization,^7^ we allow the centroid matrix to be real-valued so that each centroid vector can capture the up/down-regulated patterns of member genes in the pathways. We also incorporate pathway membership information and a GGI network into the factorization model using the framework of Bayesian learning through structured spike-and-slab priors.^3^ We also have presented efficient variational update rules for the posterior distributions. We have shown the usefulness of our methods on the synthetic and the gastric cancer data sets. To get more complete understanding of molecular biology processes, it is necessary to integrate multiple types of genomic data, copy number alternation and gene expression data, miRNA and etc. We believe that our Bayesian modeling can provide an efficient tool to implement this idea.

## Supplementary material

This supplementary material provides more details about the experiment results and the proposed method in the main text. The codes and further information are also available at https://github.com/parks-cs-ccf/BayesianSNMTF.

### TCGA gastric cancer and metastatic gastric cancer immunotherapy clinical-trial datasets: additional information

We first include the list of the selected pathways from both data sets in the experimental results section in the main text. Please see Table 2 for the TCGA gastric cancer data and Table 3 for the metastatic gastric cancer immunotherapy clinical-trial data. Further details of pathway analysis are available at our GitHub page.

**Table 2.** Summary of the top 3-ranked pathways associated with the molecular sub-types obtained from the TCGA gastric cancer dataset.

| sub-types | rank | #members | member genes |
| --- | --- | --- | --- |
| CIN | 1 | 12 | ADAMTS4,CELA1,CTRB1,DERL1,DERL2,DERL3,KLK5,MFI2,MMP26,PRSS1,PRSS3,SERPINA1 |
|  | 2 | 14 | COL2A1,COL3A1,COL9A1,COL9A2,COL9A3,COMP,FN1,MAG,MAP1B,MBP,NGFR,PLP1,PRNP,RTN4R |
|  | 3 | 13 | BARD1,BRCA1,CSTF1,CSTF2,CSTF3,FEZ1,HTATSF1,IKBKAP,MED21,PIN1,POLR2A,RBBP8,SUPT5H |
| EBV | 1 | 12 | ADAMTS4,CELA1,CTRB1,DERL1,DERL2,DERL3,KLK5,MFI2,MMP26,PRSS1,PRSS3,SERPINA1 |
|  | 2 | 13 | C3,F2,F2RL3,FCER2,HP,ICAM2,ICAM4,ITGAM,ITGAX,ITGB2,JAM2,JAM3,TJP1 |
|  | 3 | 14 | CD44,EED,FN1,ICAM4,ITGA4,ITGAE,ITGB1,ITGB7,LGALS8,MADCAM1,PXN,TLN1,VCAM1,VCAN |
| GS | 1 | 10 | CALM1,CPE,GCG,GLP1R,GPRASP1,GRM5,MEP1A,MEP1B,OPRM1,VIPR1 |
|  | 2 | 1 | GNAQ |
|  | 3 | 2 | CD200,CD200R1 |
| MSI | 1 | 3 | CCDC67,CCDC85B,EIF3E |
|  | 2 | 1 | GNAQ |
|  | 3 | 3 | NUP155,NUPL2,ZFYVE9 |

**Table 3.** Summary of the top 3-ranked pathways associated with the treatment response obtained from the metastatic gastric cancer immunotherapy clinical-trial dataset.

| sub-types | rank | #members | member genes |
| --- | --- | --- | --- |
| responder | 1 | 14 | ATN1,ECM1,ELN,FBLN1,FBLN2,FBN1,FBN2,FN1,HSPG2,ITGB1,LTBP1,MFAP2,PRELP,VCAN |
|  | 2 | 11 | CCL19,CCL21,CCL5,CCR3,CXCL11,CXCL13,CXCL9,DPP4,IGFBP7,PF4,VCAN |
|  | 3 | 10 | CCL11,CCL5,CCR3,CPAMD8,CXCL11,CXCL13,CXCL9,DPP4,FAP,PF4 |
| non-responder | 1 | 3 | SLC1A4,SLC1A5,TBC1D17 |
|  | 2 | 3 | CCDC85B,KRTAP4-12,LMO2 |
|  | 3 | 3 | NMU,NMUR1,NMUR2 |

#### More details about the proposed method

We first summarize the probability distributions used in our main paper in the following table.

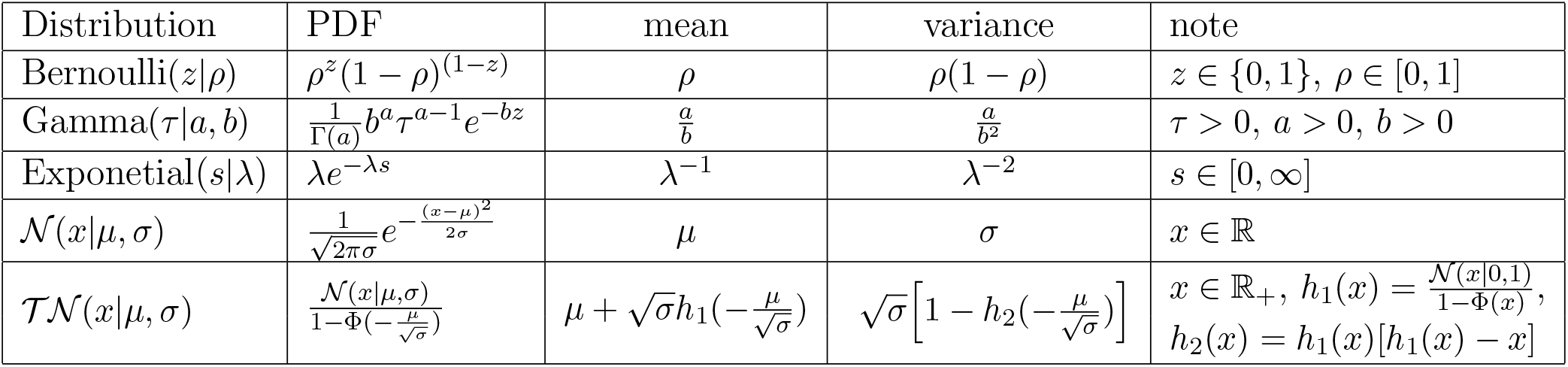

### S.1. Model Summary

The observation matrix is decomposed into the sub-matrices in the following way:

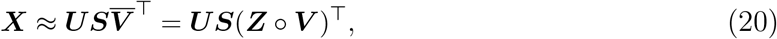

where ∘ stands for an element-wise multiplication operator. Denoting all the latent variables by Θ ≜ {***S, V, Z, G***}, the joint probability distributions of the model is given as follows:

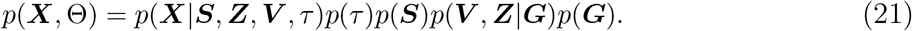

where

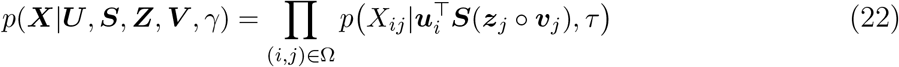

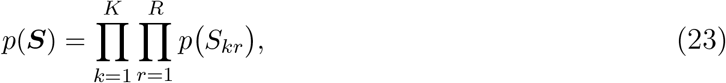

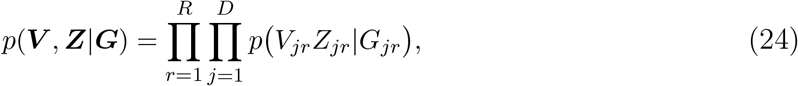

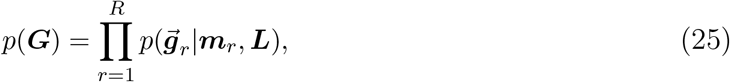

where 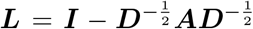 is a normalized Laplacian matrix and ***A*** is an adjacency matrix driven from a protein-protein interaction network (*A_ij_* = 1 if *i* ≠ *j* and there is a connection between the genes *i* and *j* on the network, and otherwise *A_ij_* = 0). Note that, the mean vector ***m***_*r*_ is set according to the membership information encoded in the pathways *Z*^0^: *m_jr_* = ***ξ***_+_ if 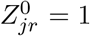, otherwise *m_jr_* = *ξ*_−_, where *ξ*_+_ > 0 and *ξ*_−_ < 0 (in our all experiments, we use *ξ*_+_ = 3 and *ξ*_−_ = −5). The form of each probability distribution in (21) is given as follows:

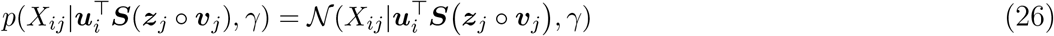

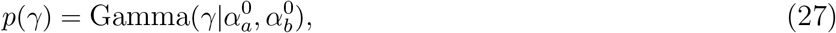

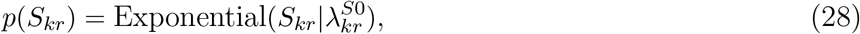

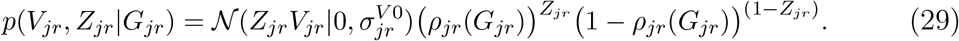

### S.2. Variational Inference

The posterior distributions over the latent variables are approximately computed in the framework of variational inference. The variational distributions that approximate the true posterior distributions over the latent variables are assumed to be factorized as follows:

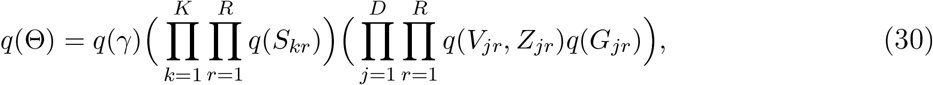

where

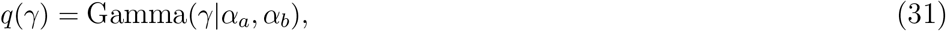

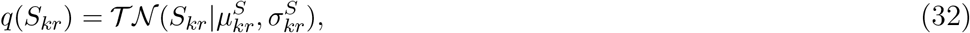

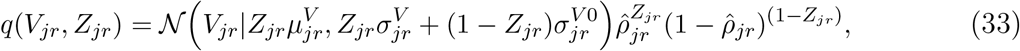

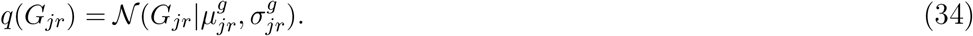

The variational distributions can be computed by maximizing the lower bound with respect to (w.r.t.) the variational distributions. Denoting a set of all the latent variables by Θ = {*γ*, ***S, Z, V, G***}, we can show that the log-likelihood can be decomposed as follows:

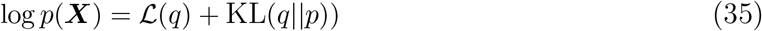

where

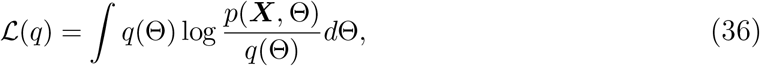

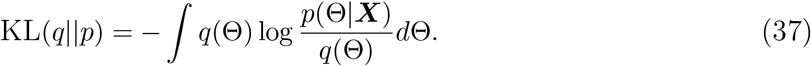

where KL(*q*||*p*) is Kullback-Leibler (KL) divergence between the variational distribution and the true posterior distribution and is always nonnegative (KL(*q*||*p*) =0 if and only if *q* = *p*). Thus, we can easily see that the log-likehood is lower-bound by the variational lower bound 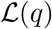 and thus the variational distributions can be updated by maximizing 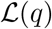 w.r.t. their parameters. Note that, the variational bound of our model is expressed as follows:

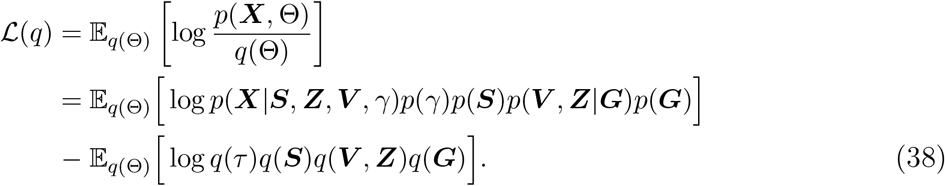

The variational distributions *q*(*γ*), {*q*(*S_kr_*)} and {*q*(*V_jr_, Z_jr_*)} can be updated in closed from. Letting Θ_*l*_ be the variable we want to update at each turn and Θ\^*l*^ be the remaining variables, the optimal solution of *q*(Θ_*l*_) can be given by the stationary condition for the factor *q*(Θ_*l*_) in the maximization problem, i.e., 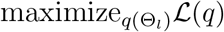:

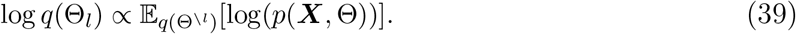

On the other hand, for {*q*(*G_jr_*)}, their means and variances can be updated by any iterative gradient-based optimization methods, e.g., limited-memory BFGS used in our experiments. We provide detailed derivations of each update in the following subsections.

#### S.2.1. Update of the variational distributions over γ and S

The variational distribution over the precision *γ* can be updated as follows:

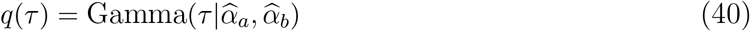

where

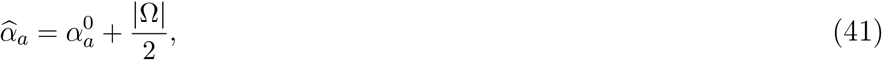

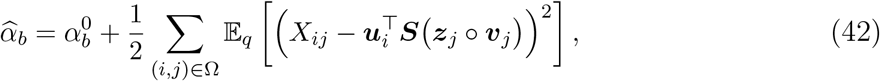

where Ω is a set of indices of the observations and

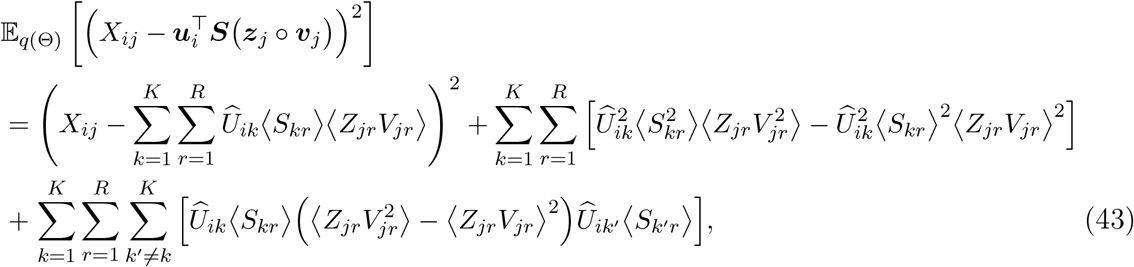

where 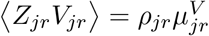 and 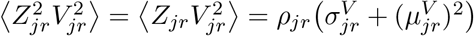.

The variable ***S*** can be updated as follows:

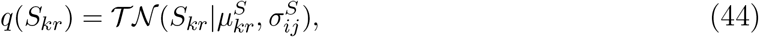

where

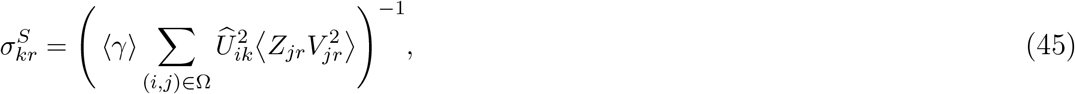

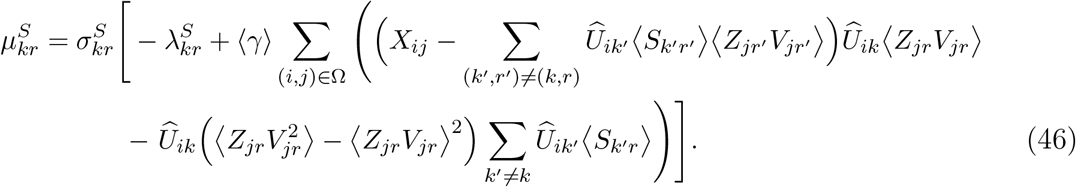

#### S.2.2. Update of the variational distributions over Z and V

Each pair of elements, {*Z_jr_, V_jr_*}, can be updated by the inference method in.^16^ From the stationary condition for *q*(*Z_jr_, V_jr_*) when maximizing the variational bound 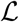 in (38), we have

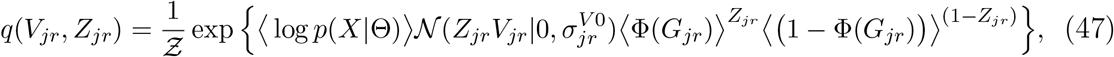

where 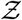 is a normalization constant. We can see that *q*(*V_jr_, Z_jr_*) can be factorized as

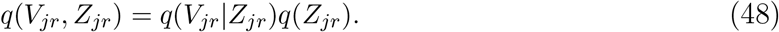

The marginal probability distribution over the binary variable *Z_jr_* can be calculated as follows:

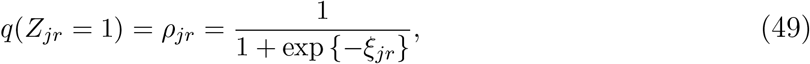

where

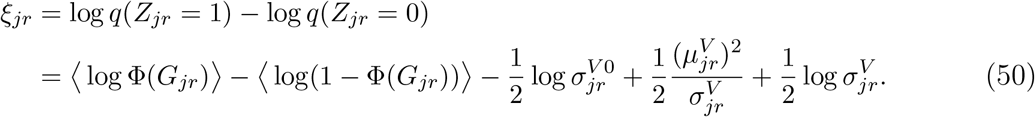

where the expectations in the second equality are approximated using Jensen’s inequality:

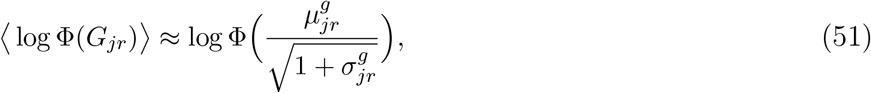

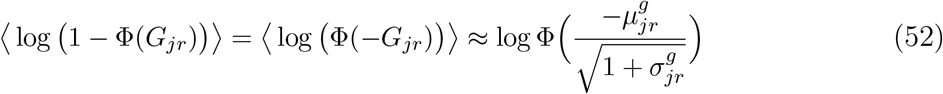

The conditional variational distribution of *V_jr_* given *Z_jr_* is given by

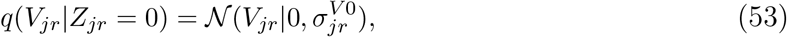

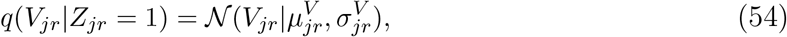

where

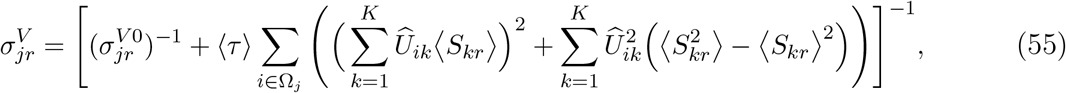

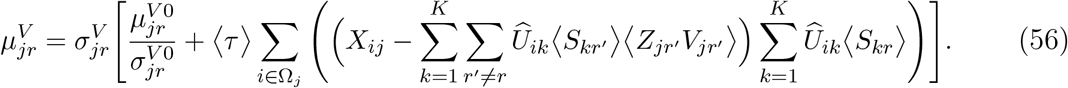

As a summary, the joint probability distribution is simply rewritten as follows:

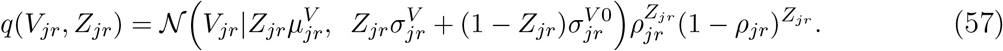

#### S.2.3. Update of the variational distributions over G

The optimization problem (38) can be reduced as follows:

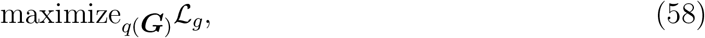

where 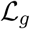 is a function including only terms which are related to the variable ***G***:

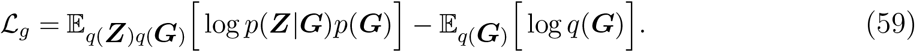

The first term of 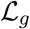 in eq. (58) can be calculated as follows:

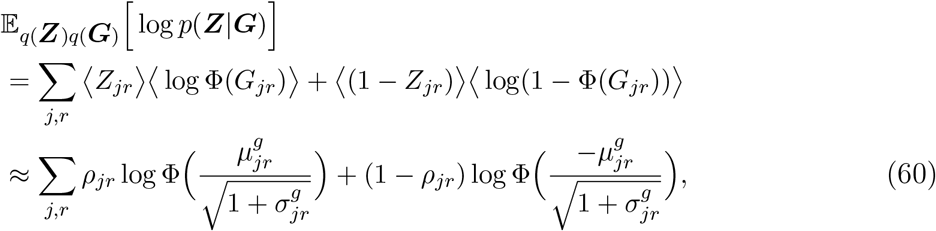

where we have used the same techniques (using Jensen’s inequality) as in the previous subsection. We then calculate the third term, a sum of entropy terms of *R* Gaussian distributions:

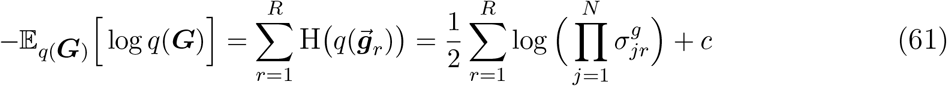

where *c* is a constant, which is independent of the variable ***G***. The second term is a cross entropy between two Gaussian distributions, *p*(*G*) and *q*(*G*), calculated as follows:

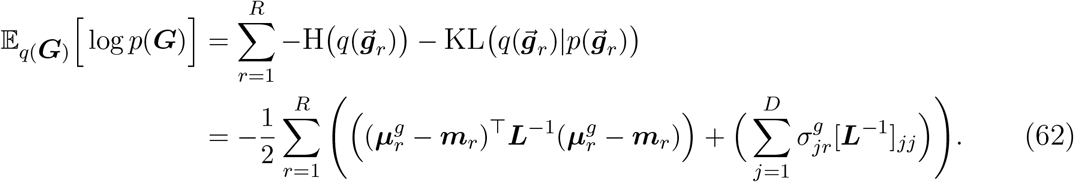

The gradient of 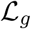 w.r.t. the parameters 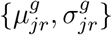 also can be easily calculated. We update these parameters using limited-memory BFGS in our experiments

### S.3. Experimental settings

We here explain how to initialize our factorization model, specifically the parameters of the prior distributions (referred to as prior hyperparameters) and those of the variational distributions (referred to as variational parameters). Regarding the variational parameters (e.g., the mean and variance for the case where the variational distribution is Gaussian) note that the variational distributions are updated cyclically, i.e., for each step, we update one variational distributions, fixing the others. Therefore, we need to initialize some (though not all) variational parameters as well as the prior hyperparameters.

The prior hyperparameters are set as follows:

- (For the noise precision *γ*) 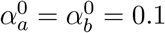
- (For each association *S_kr_*) 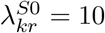 for all *k, r*
- (For each element in the centroid, *V_jr_*) 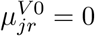 and 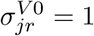
- (For the prior mean vectors of the GPs, ***m***_*r*_) *ξ*_+_ = 5 and *ξ*_−_ = −5

Based on experience from experimentation on synthetic and various gene expression datasets, the factorization results of the method are generally not sensitive to the most parameters’ initial settings. However, we should note that *ξ*_+_ and *ξ*_−_ represent the prior belief in the initial pathway membership information ***Z***^0^. If we set *ξ*_+_ = *ξ*_−_ = 0, the prior probability of the on-off binary variable *Z_jr_* = 1 is 0.5 regardless of whether the *r*th pathway includes the *j*th gene or not, i.e., for both cases 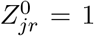 and 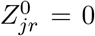. The more extreme values *ξ*_+_ and *ξ*_−_ have, i.e. *ξ*_+_ >> 0 and *ξ*_−_ << 0, the stronger the prior belief we place on the initial pathway information ***Z***^0^. The setting we use in our experiments (*ξ*_+_ = 5 and *ξ*_−_ = −5) usually gives satisfactory factorization results. However, users can adjust them according to their prior belief in the pathway information.

The variational parameters are set as follows:

- (For *S_kr_*) 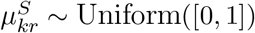, for all *k, r*
- (For ***V***) each column is set to the centroid from *K*-means ran on the input data ***X*** if *R* < *N* and to the randomly chosen sample (row) from the input matrix ***X*** otherwise.
- (For *G_jr_*) 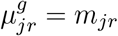 and 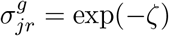, where 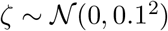

### S.4. Bayesian semi-nonnegative- vs Point estimate non-negative factorization

As explained in the main paper, many types of genomic data are given in a form of a real-valued matrix after relevant normalization or transformation steps. However, the NMF formulation does not permit negative values in the inputted observation matrix. Thus, one of the standard ways to handle negative values for the NMF formulation is to fold the original matrix by columns:^4^ every column (gene) will be represented in two new columns in a new observation matrix, one of which contains only the positive values (upregulations) and the other column only the magnitudes of the negative values (downregulations). However, the folding approach incurs increased computational complexities: the number of columns in the input matrix is doubled, and the gene-gene interaction network is 2^2^ times larger. On the other hand, our approach (motivated by semi-nonnegative factorization^7^) allows the centroid matrix to have negative values but still imposes nonnegativity constraints on the encoding matrix. Furthermore, we implement our semi-nonnegative tri-matrix factorization algorithm in the framework of Bayesian learning. In the following subsections, we provide two specific examples that show the superiority of our method over non-negative tri-matrix factorization (NTriPath^2^) which is implemented using the folding approach to deal with negative values in the input matrix.

Before presenting the details of the simulated experiments, we first provide a brief introduction to NTriPath. The objective of the method is again to approximate the input matrix ***X*** as a product of the three small matrices, ***U*** (the sub-type indicator matrix), ***S*** (the association matrix) and 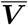 (the centroid matrix), i.e., 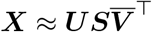. With ***U*** fixed, ***S*** and 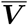 are estimated by minimizing the following objective function under the non-negativity constraints:

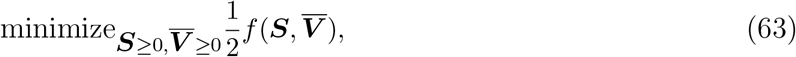

where the objective function is define as:

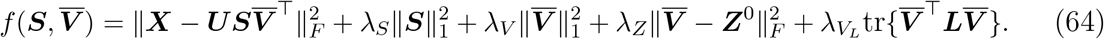

The matrices ***S*** and 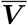 are updated by multiplicative rules to ensure the non-negativity constraints. ^2^ Note that NTriPath involves 4 regularization parameters which should be specified by the user. Identification of associations between sub-types and pathways from input data is clearly an unsupervised learning problem since true associations generally are unknown. Therefore, it is unclear how to tune the regularization parameters of NTriPath for the given input data. In addition, the method incurs a high computational burden for large scale datasets when the best combination of the hyperparameters is searched among a set of candidate values (the search space being a 4D grid space) by cross validation. For simplicity, we fix λ*_V_L__* = λ_*Z*_ = 1 as in our previous work. We tune only λ_*S*_ and λ_*V*_ which are related to the sparseness of the metrics. We select the best regularization constants from 2D grid space (the grid space along each dimension being defined as {0.001, 0.005,0.01,0.05,0.1, 0.5,1}) by finding the combination which gives the least reconstruction error. Meanwhile, our Bayesian method is able to automatically tune model complexity, including the noise precision, by integrating over all the latent variables.

#### S.4.1. Baseline example

We begin by presenting a simple example that contains a basic structure in the observation matrix and other inputs. We discuss the results of this straightforward settings before examining the two cases in which our proposed Bayesian framework provides clear advantages over the folding approach. A detailed overview of data generation is now presented for our first example.

Inspired by the biological setting of the gene expression application in the main paper, we use the same terminology here as in the main script (i.e. subgroups, genes, pathways) in discussing the problem formulation and results of our simulated experiments. As in the main application, these experiments attempt to decompose patterns of upregulated and down regulated genes within different patient subgroup. In this preliminary example, as well as in subsequent experiments, the observation matrix ***X*** ∈ ℝ^200×800^ consists of 4 subgroups (each containing 50 samples) with some defined pattern among the 800 genes (which are grouped into sets of 100). Within each subgroup, a set of 100 genes can represent upregulation, down regulation or background noise. For upregulated and downregulated genes, samples are drawn from a Gaussian 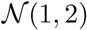 or Gaussian 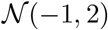, respectively. For background noise, samples are drawn from a Gaussian 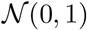. A simple block structure was determined with each subsample containing 2-3 “selected” (either upregulated or downregulated) gene sets (Fig. 4a). Subgroups are encoded in 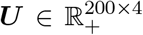 using simple 1-of-*K* encoding (*U_ij_* ∈ {0,1} and Σ_*j*_*U_ij_* = 1) (Fig. 4b). Gene-pathway prior knowledge is encoded in the matrix 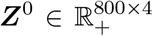 and is initialized to contain similar structure to the observation ***X***. That is, we allow ***Z***^0^ to contain 4 pathways that reflect the same pattern of selected genes within the subgroups of ***X*** by setting 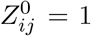 for all the genes *i* that are upregulated or downregulated within the subgroup that we have designated to pathway *j* (Fig. 4c). The gene-gene interaction network ***A*** ∈ ℝ^800×800^ is initialized with approximately 10% sparcity and contains random symmetric connections (with no self connections; Fig. 4d). Our motivation for this simple design is to allow our method to arrive at a simple and predictable solution for the model’s learned factors, namely, the subgroup-pathway association matrix 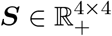, the real-valued pathway-gene association matrix ***V*** ∈ ℝ^800×4^ and the updated pathway-gene binary membership matrix 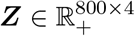.

Fig. 5 and Fig. 6 show the factorization results of our method and NTriPath, respectively. Both methods produce correct estimates of the true association matrix ***S*** although our method’s estimate is more clearly separable (Fig. 5a).

**Fig. 4.**
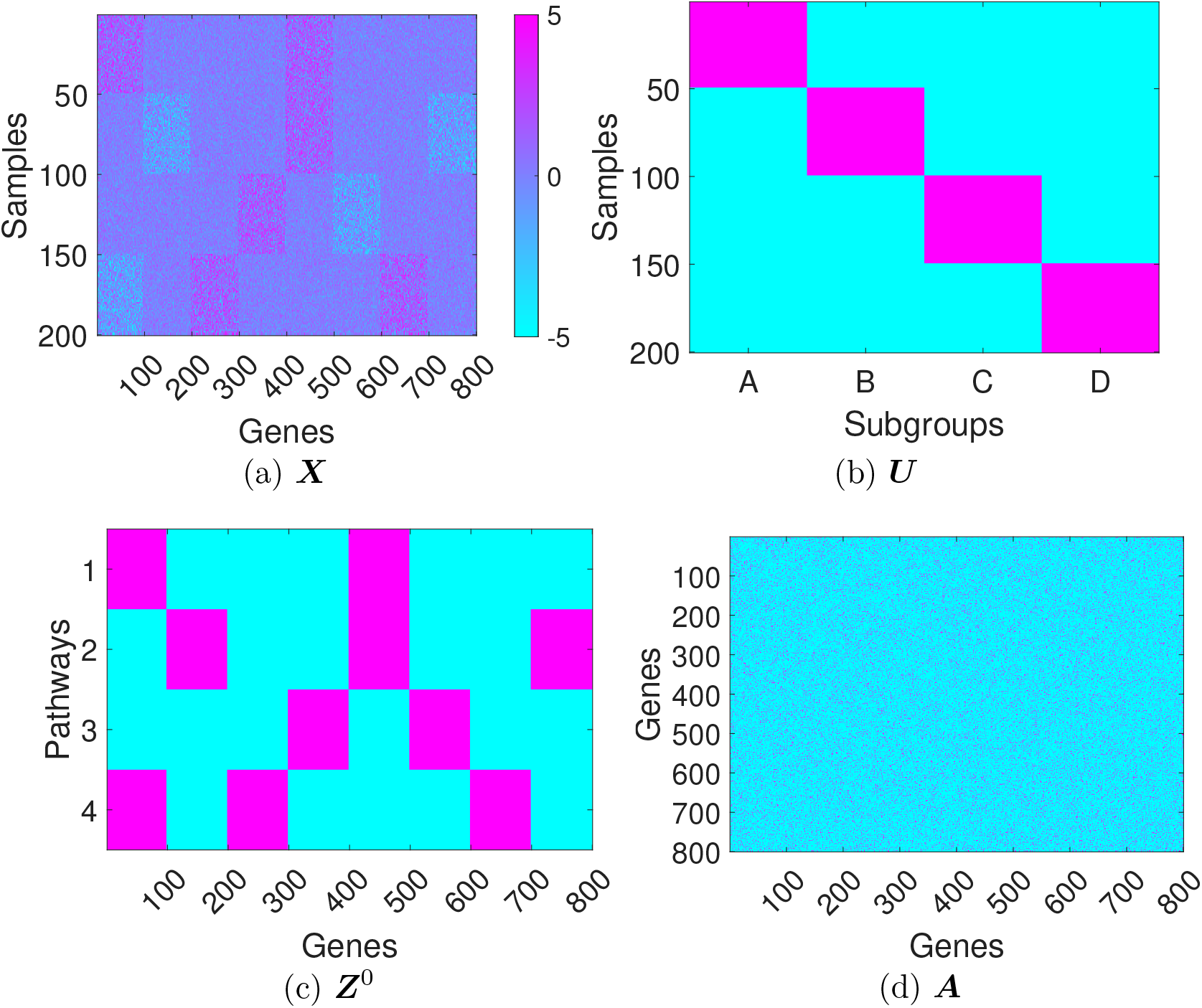
Inputs to both algorithms.

**Fig. 5.**
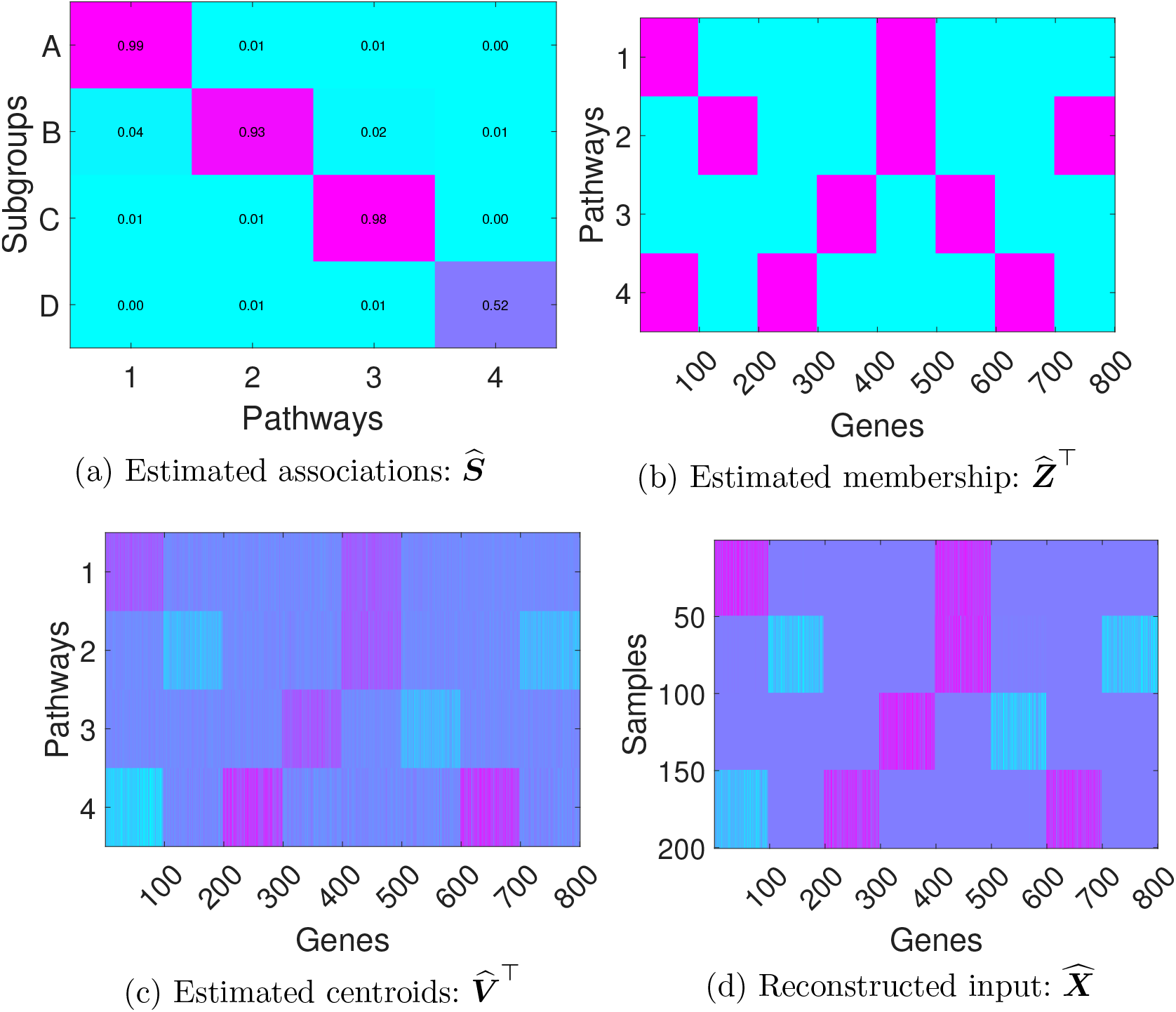
Factorization result from our method.

**Fig. 6.**
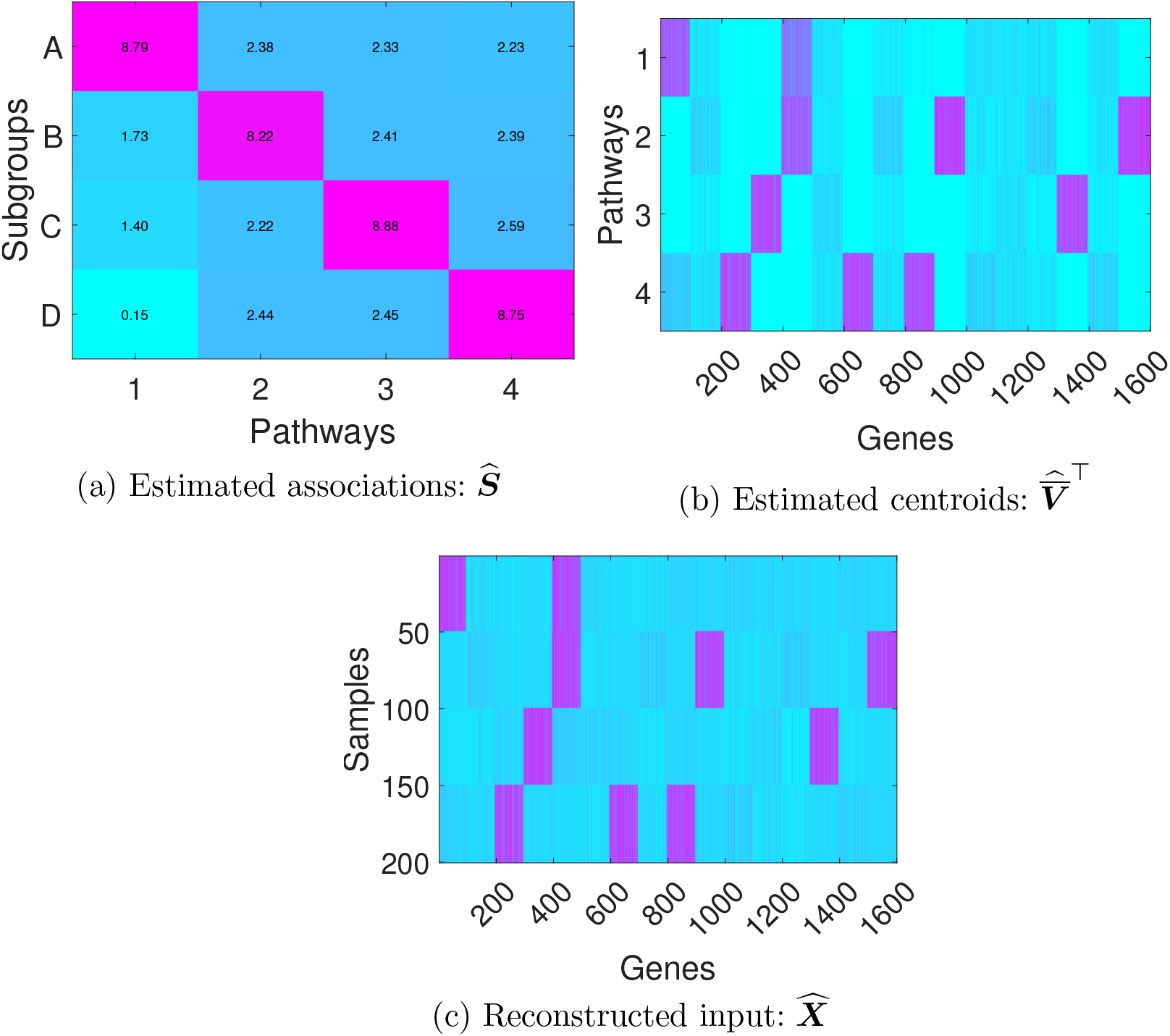
Factorization results from NTriPath.

#### S.4.2. Limitations of NMF methods based on the folding approach

We here provide a simple example where NTriPath, employing the folding approach to deal with negative values, fails to correctly estimate associations between sub-types and pathways from a real-valued input matrix. The main reason for this incorrectly learned association matrix is that the folding approach breaks the original underlying patterns of the input matrix by separating non-negative and negative values. In fact, this issue is problematic not only for NTriPath but for all NMF methods that are based on the folding approach. Note that, however, our factorization method is free from this issue due to the semi-nonnegative modeling, which is one of the primary advantages of our method compared to NTriPath and other NMF based methods using the folding approach.

Fig. 7 shows how the input matrix ***X*** is generated based on the baseline example in the previous subsection (Fig. 7a) and how the new non-negative input matrix ***X***_*new*_ is constructed by the folding approach i.e., *X*_new_ ≜ [max(*X*, 0), max(−*X*, 0)] (Fig. 7b). We assume that expression values at the noisy block of genes in the first sub-type samples are drawn from i.i.d. Gaussian distributions (white Gaussian noise) with a high variance, i.e., 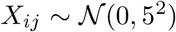. In other words, this data block (represented as *X*[1: 50,101: 200] in MATLAB language) contains just random noise values. However, these negative and non-negative noisy elements become strong signal blocks when the input matrix is transformed by the folding approach (see *X_new_*[1: 50,101: 200] and *X_new_*[1: 50,901: 1000] in Fig. 7b). Thus, NTriPath tries to fit both noisy blocks, which should be ignored as noise, by adjusting the association matrix ***S*** and other factorization results. Fig. 8a clearly supports this discussion. We can see that the estimated association matrix 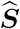 from NTriPath fails to recover the expected subgroup-pathway associations and thus the top pathways associated with each sub-type are also incorrect. However, as mentioned before, our factorization method based on the semi-nonnegative modelling yields the correct estimate for associations without being affected by the presence of the noise block (Fig. 8b).

**Fig. 7.**
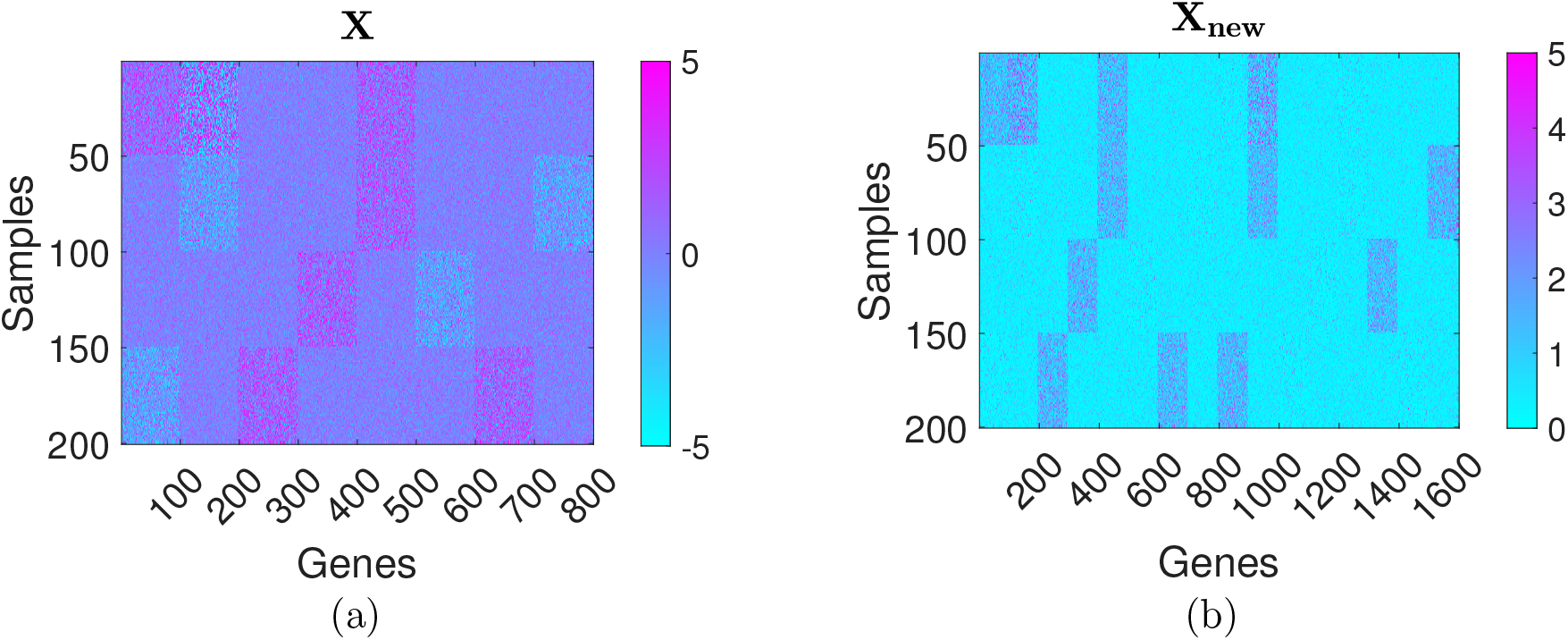
A simple example where NMF methods based on the folding approach fail to correctly estimate true association between sub-types and pathway from a real-valued input matrix: a) There is a noisy block in the original input matrix, i.e., *X*[1: 50,101: 200]; b) the negative and non-negative values in this block become strong signals after transformed by the folding approach.

**Fig. 8.**
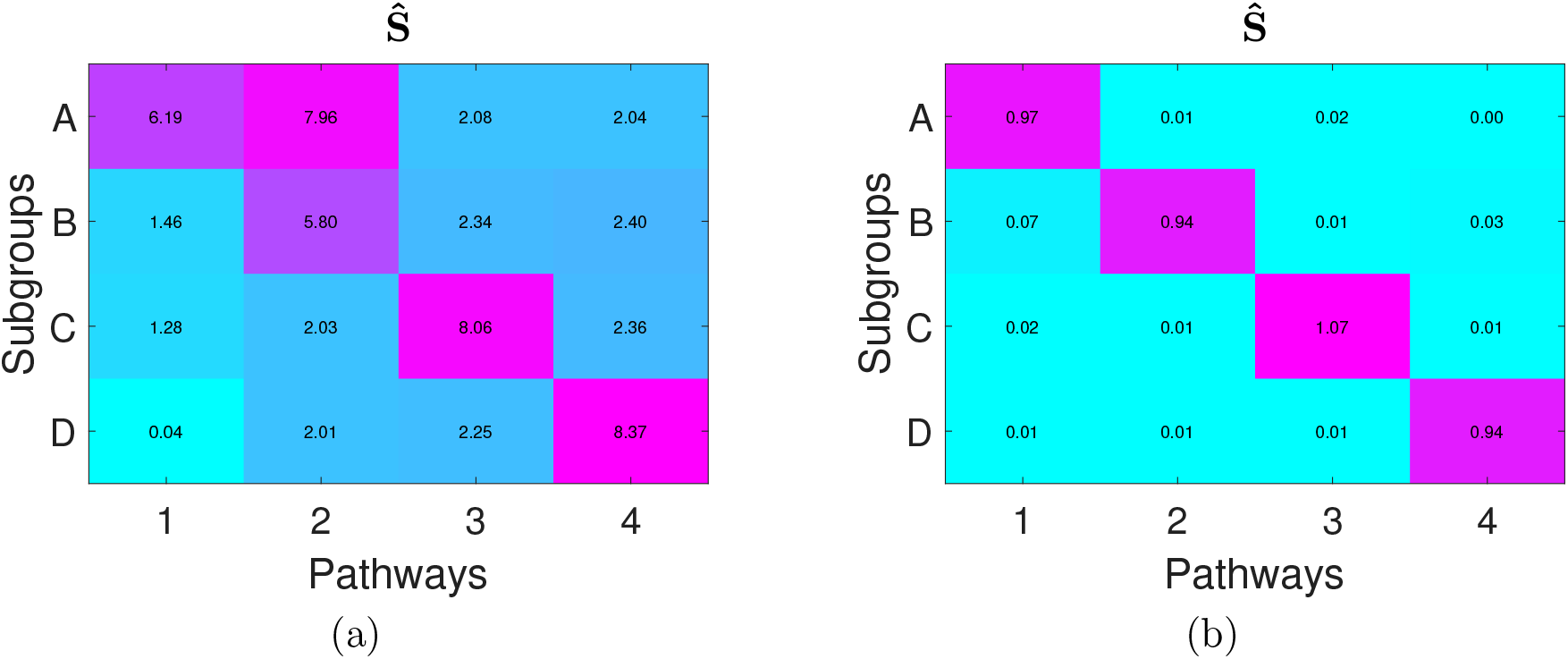
The estimated association matrices from (a) NTriPath and from (b) our method. The presence of the noisy block causes NTriPath to make an incorrect association identification.

#### S.4.3. Robustness against noise

We compare the performance of our Bayesian factorization method and of NTriPath in the case where the input matrix is contaminated by background noise of different noise levels. Our objective here is to show whether each method is robust against increasing noise. We assume that the observation matrix 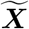 is generated by adding white Gaussian noises to the data input matrix ***X*** defined in the previous subsection, *Baseline example*, i.e., 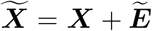, where 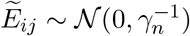 and the noise variances 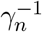 increases incrementally from 1^2^ to 10^2^. We train both methods on the noisy observation matrix 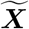 to test how each method performs in correctly identifying the associations between the sub-types and the pathways.

We report the performance of both methods in Fig. 9. Since we know the ground truth associations for this dataset, we can calculate the accuracy of each method based on how many associations each method correctly predicts. We repeat each experiment 20 times at each noise variance. As we can see in the figure, our method shows overall stable performance in the entire range of the noise variance. Note that our method shows the slightly worse performance at the first three noise levels but maintains almost the same performance as the noise level increases. We hypothesize that the sub-optimal performance of our method at the first three noise levels is caused by improperly randomized initialization points. However, our Bayesian factorization method still works well at high noise levels, where the performance of NTriPath dramatically diminishes. This supports our claim of the greater robustness of our method against noise relative to NTriPath.

**Fig. 9.**
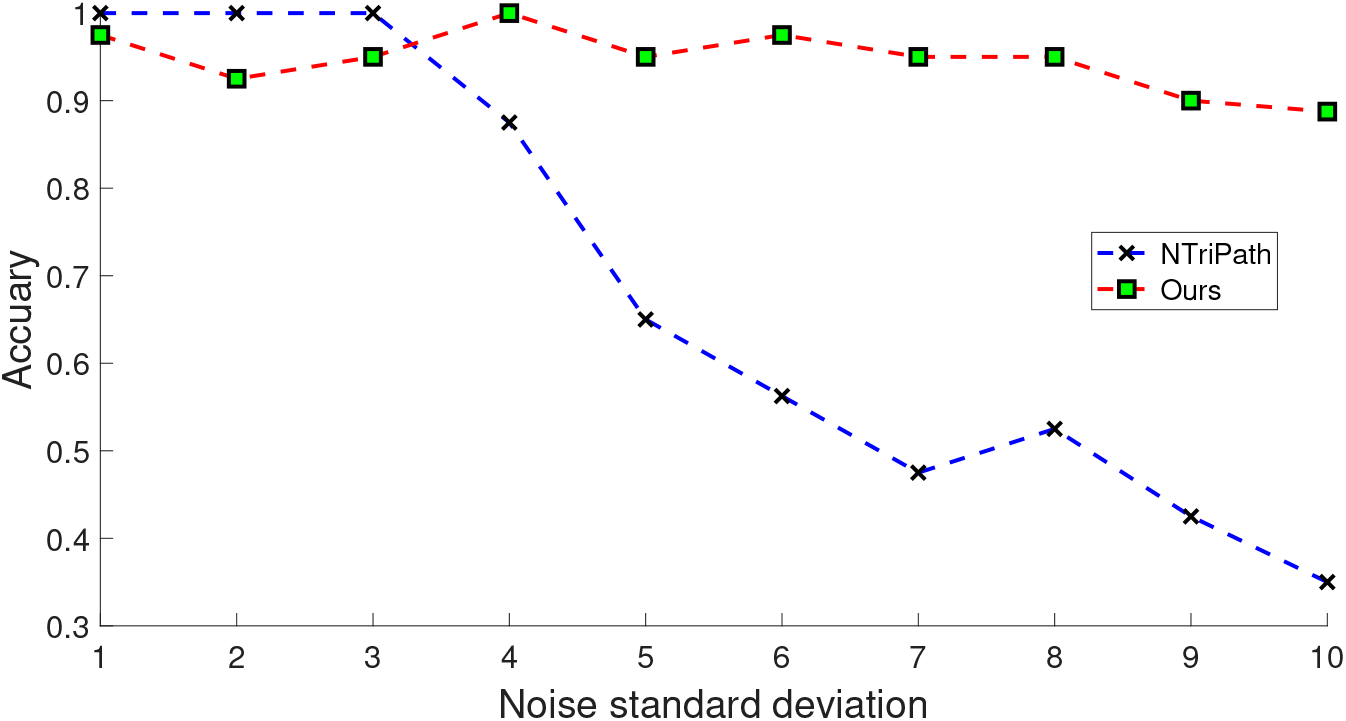
Robustness of both methods against noise: the prediction performance of our method is compared to NTriPath as the noise variance increases.

## Footnotes

* The data was downloaded from CBioportal (http://www.cbioportal.org/). The downloading option was ‘TCGA stad rna seq v2 mrna’ (RNASeq V2 RSEM normalized expression values).

† https://thebiogrid.org/. The version is BIOGRID-ORGANISM-Homo sapiens-3.4.153.

## References

1. C. Ding, T. Li, W. Peng and H. Park, Orthogonal nonnegative matrix tri-factorizations for clustering, in Proceedings of the ACM SIGKDD Conference on Knowledge Discovery and Data Mining (KDD), (Philadelphia, PA, 2006).

2. S. Park, S.-J. Kim, D. Yu, S. Pea-Llopis, J. Gao, J. S. Park, B. Chen, J. Norris, X. Wang, M. Chen, M. Kim, J. Yong, Z. Wardak, K. Choe, M. Story, T. Starr, J.-H. Cheong and T. H. Hwang, An integrative somatic mutation analysis to identify pathways linked with survival outcomes across 19 cancer types, Bioinformatics 32, 1643 (2016).

3. M. R. Andersen, O. Winther and L. K. Hansen, Bayesian inference for structured spike and slab priors, in Advances in Neural Information Processing Systems (NIPS), eds. Z. Ghahramani, M. Welling, C. Cortes, N. D. Lawrence and K. Q. Weinberger 2014 pp. 1745–1753.

4. P. Kim and B. Tidor, B. subsystem identification through dimensionality reduction of large-scale gene expression dat, Genome Research 13, p. 17061718 (2003).

5. S. T. Kim, R. Cristescu, A. J. Bass, K.-M. Kim, J. I. Odegaard, K. Kim, X. Q. Liu, X. Sher, H. Jung, M. Lee, S. Lee, S. H. Park, J. O. Park, Y. S. Park, H. Y. Lim, H. Lee, M. Choi, A. Talasaz, P. S. Kang, J. Cheng, A. Loboda, J. Lee and W. K. Kang, Comprehensive molecular characterization of clinical responses to pd-1 inhibition in metastatic gastric cancer, Nature medicine 24, p. 14491458 (2018).

6. K. Devarajan, Nonnegative matrix factorization: An analytical and interpretive tool in computational biology, PLoS Computational Biology 4 (2008).

7. C. Ding, T. Li and M. I. Jordan, Convex and Semi-Nonnegative Matrix Factorizations, Tech. Rep. 60428, Lawrence Berkeley National Lab (2006).

8. C. M. Bishop, Pattern Recognition and Machine Learning (Information Science and Statistics) (Springer-Verlag, Berlin, Heidelberg, 2006).

9. T. Brouwer and P. Lio’, Fast bayesian non-negative matrix factorisation and tri-factorisation, in NIPS 2016 Workshop: Advances in Approximate Bayesian Inference, 2016.

10. R. Cristescu, J. Lee, M. Nebozhyn, K.-M. Kim, J. Ting, S. S. Wong, J. Liu, Y. Gang Yue, J. Wang, K. Yu, X. Ye, I.-G. Do, S. Liu, L. Gong, J. Fu, J. Gang Jin, M.-G. Choi, T. Sung Sohn, J. Ho Lee and A. Aggarwal, Molecular analysis of gastric cancer identifies subtypes associated with distinct clinical outcomes, Nature medicine 21 (04 2015).

11. B. H. Sohn, J.-E. Hwang, H.-J. Jang, H.-S. Lee, S. C. Oh, J.-J. Shim, K.-W. Lee, E. H. Kim, S. Y. Yim, S. H. Lee, J.-H. Cheong, W. Jeong, J. Y. Cho, J. Kim, J. Chae, J. Lee, W. K. Kang, S. Kim, S. H. Noh, J. A. Ajani and J.-S. Lee, Clinical significance of four molecular subtypes of gastric cancer identified by the cancer genome atlas project, Clinical Cancer Research (2017).

12. S. Suthram, J. T. Dudley, A. P. Chiang, R. Chen, T. J. Hastie and A. J. Butte, Network-based elucidation of human disease similarities reveals common functional modules enriched for pluripotent drug targets, PLoS Comput Biol 6, p. e1000662 (2010).

13. K. Ganesh and J. Massague, TGF-Inhibition and Immunotherapy: Checkmate, Immunity 48, 626 (04 2018).

14. L. L. van der Woude, M. A. J. Gorris, A. Halilovic, C. G. Figdor and I. J. M. de Vries, Migrating into the Tumor: a Roadmap for T Cells, Trends Cancer 3, 797 (11 2017).

15. S. Monti, P. Tamayo, J. P. Mesirov and T. R. Golub, Consensus clustering: A resampling-based method for class discovery and visualization of gene expression microarray data., Machine Learning 52, 91 (2003).

16. M. K. Titsias and M. Lázaro-Gredilla, Spike and slab variational inference for multi-task and multiple kernel learning, in Advances in Neural Information Processing Systems (NIPS), eds. J. Shawe-Taylor, R. S. Zemel, P. L. Bartlett, F. Pereira and K. Q. Weinberger 2011 pp. 2339–2347.

